# A unique sixth-order reversible assembly of hexameric mimetic nucleic acid

**DOI:** 10.64898/2026.11.12.766261

**Authors:** Sajena Kanangat Saraswathy, Jonathan B. Chaires, Gary B. Schuster, Nicholas V. Hud

## Abstract

Complementary dipeptides bearing the non-canonical nucleobases melamine and cyanuric-acid (denoted as α-DPNA) assemble into a cylindrical hexameric structure containing a stacked heterocyclic core with its β-sheet peptide backbones on the surface. Circular dichroism spectroscopy and singular-value decomposition show that the evolution of CD spectra after mixing equimolar strands is an apparent two-state process, accounted for by only the spectra of free strands and the assembled hexamer, with no detectable intermediates. Equilibrium measurements reveal a highly cooperative, enthalpically driven assembly with a molecularity of 6.2 ± 0.2 and an association constant of (2.3 ± 0.2) × 10^22^ M^−5^. Over total strand concentrations of 20–100 µM, the assembly progress curves are described by an apparent reversible sixth-order macroscopic rate law. This deceptively simple behavior cannot represent an elementary six-strand collision. The apparent forward rate constant has a negative activation energy, showing that the observed rate law masks additional steps in the assembly reaction. A minimal model in which strand pre-isomerization precedes hexamer formation restores a physically reasonable positive activation energy while retaining the observed two-state CD behavior. Constrained simulations further show that a sequential pathway with low-population, spectroscopically silent, intermediates can give rise to the same deceptively simple sixth-order progress curve. These results provide an unusual example of highly cooperative hexamer formation and show how a simple high-order macroscopic rate law can conceal a more complex microscopic assembly mechanism. The thermodynamics and kinetics provide insights into how and why the hexamer assembles, offering some design principles for supramolecular assemblies in water.

**Table of Contents Graphic:** 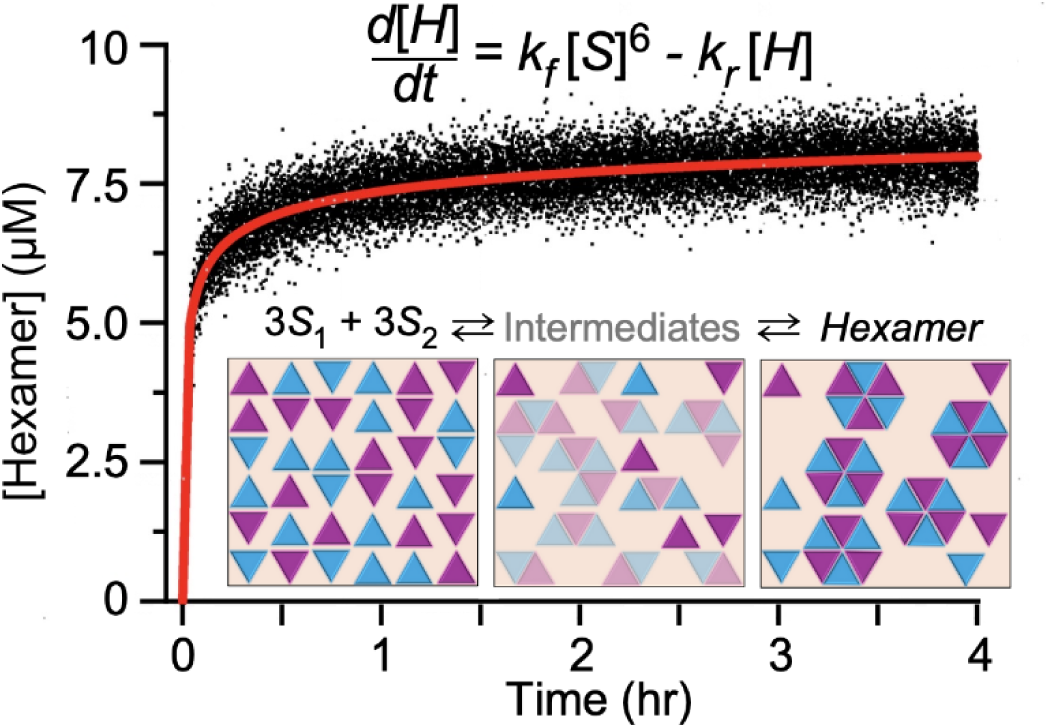

## Introduction

Water-soluble supramolecular polymers of paired and stacked heterocycles are of interest for the development of novel materials and have been proposed as possible ancestral forms of RNA^1^. The non-canonical nucleobases 2,4,6-triaminopyrimidine and substituted cyanuric acid in water can, for example, self-assemble by hydrogen bond pairing and stacking interactions into stable polymeric structures that feature a planar hexameric rosette core^2^. Melamine and cyanuric acid were also found to self-assembly into more amorphous structures^3^. A variety of efforts were made to attach non-canonical bases to peptide or sugar-phosphate backbones to even better match potential primordial oligomers^3–5^. Recently this laboratory designed and synthesized homo-tetrameric strands with melamine or cyanuric acid “bases” attached to octapeptide backbones (denoted α-(M)_4_ and α-(C)_4_, respectively, to explore their interactions and assembly (Scheme 1). The complementary strands self-assembled into a stable hexameric structure featuring a cylindrical core of stacked, base-paired bases, with the peptide backbones arranged in an ordered beta-sheet configuration surrounding the core, reminiscent of a small beta barrel^6^. A model of the structure is shown in Scheme 1.

**Scheme 1.**
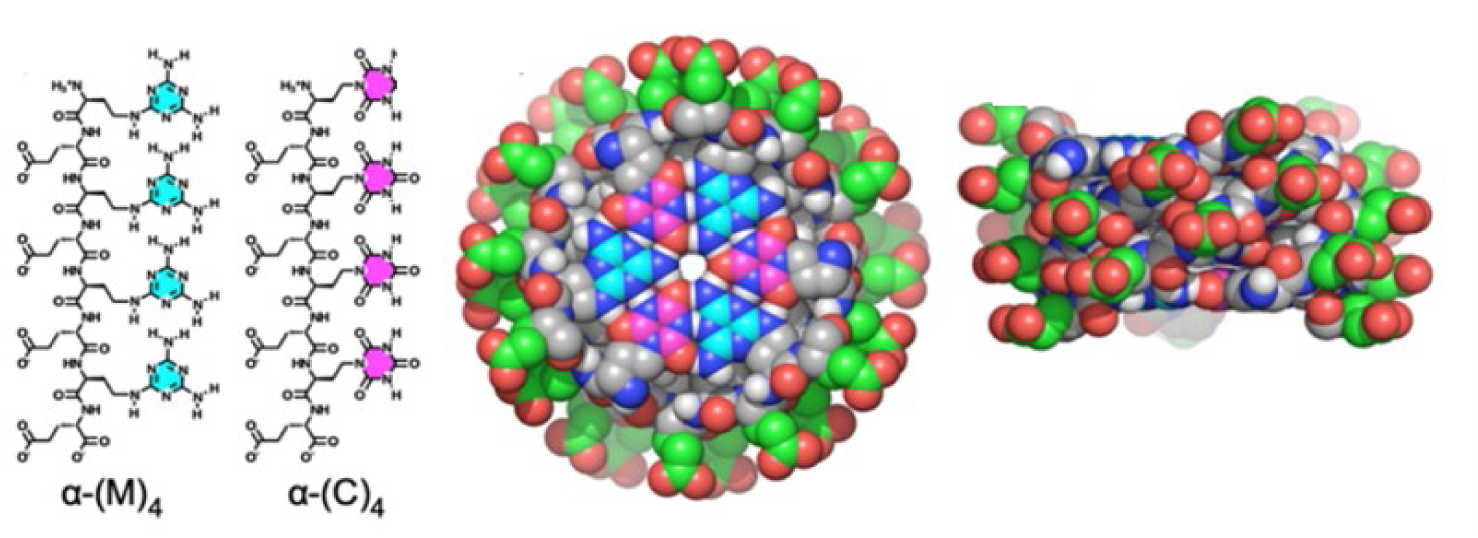
Chemical structures of α-DPNA strands (left) and two orthogonal views (right) of the hexameric assembly in space-filling representation.

Nucleic acids are remarkable in their ability to form multimolecular assemblies, with well-documented examples of the canonical duplex along with triplex^7^, tetramolecular G-quadruplex^8–12^ and the tetramolecular C-rich i-motif structures^13^. The thermodynamics and kinetics of these forms have been well-studied to define the mechanisms and driving forces for their formation^8, 10–11, 14–18^. In contrast, detailed thermodynamic and kinetic studies of the assembly of non-canonical bases and their oligomeric constructs are less common, with some exceptions^3^. Here we aim to improve this situation for the assembly of the octapeptides α-(M)_4_ and α-(C)_4_ into a hexamer.

We find that these strands assemble in an apparent single-step, reversible, sixth-order reaction. Such a reaction must be exceedingly rare. We did not know of any prior examples and an exhaustive AI-driven literature search did not produce any publications containing experimental examples. Kineticists typically dismiss the possibility of such a higher-order reaction since it would require the simultaneous collision of six reactant molecules. As early as 1938, Farrington Daniels in his classic kinetics text^19^ noted in his discussion of reaction orders that: “Third order reactions are rare because triple collisions are statistically improbable, and equations of the fourth order and higher orders do not even need to be considered.” Nonetheless, tetramolecular quadruplex formation was found to follow an apparent fourth-order rate law providing a precedent for questioning such kinetic dogma^8, 10–12^ We show here evidence for an apparent macroscopic reversible sixth-order progress curve of the octapeptides α-(M)_4_ and α-(C)_4_ into a hexamer but also show that our primary data mask hidden complexity that arises from a more complex underlying reaction, the nature of which we explore.

## Results & Discussion

### Circular dichroism provides a means to monitor strand assembly

Figure 1 shows the results of circular dichroism (CD) experiments used to monitor the assembly of equimolar mixtures of octapeptides α-(M)_4_ and α-(C)_4_. Panel A shows the initial mixing experiment at 160 µM total strand concentration intended to drive assembly to completion. These spectra are unlike any typical nucleic acid spectra that originate primarily from base stacking. Instead, these CD spectra are seen to evolve from one characteristic of a random peptide chain into a spectrum with the features of a beta sheet. These spectra comprise a matrix that can be analyzed by singular value decomposition (SVD)^20^ to evaluate its rank and complexity (Fig. S1-S2). SVD showed conclusively that the data in panel A can be accounted for by a linear combination of only two spectral species (and their difference), consistent with a single-step, two-state assembly reaction. As validation, the family of spectra can be reconstructed using only these two basis spectra and their difference (Fig. S2) to produce a family of spectra with greatly reduced noise that, when subtracted from the original data, yields residuals that are by all statistical measures random. A key consequence of this analysis is that CD can be used to quantitatively determine the concentrations of free and assembled strands from any given spectrum by the linear combination of the two properly normalized basis spectra (Fig. 1B). There are no detectable intermediates with distinctive CD signatures.

**Figure 1.**
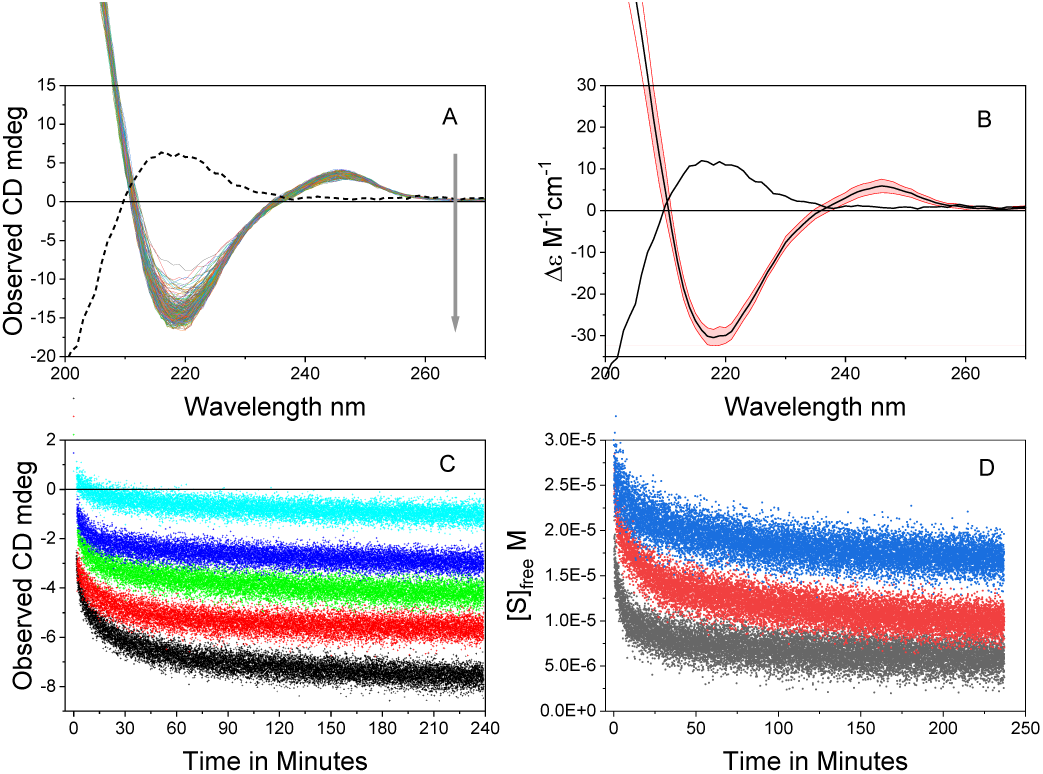
Circular dichroism of the assembly of the octapeptides α-(M)_4_ and α-(C)_4_. A) Changes in CD spectra from 2-1400 minutes after mixing equimolar strands to a final total strand concentration of 160 µM. The dashed line shows the average spectrum of the unassembled strands at this concentration. B) Molar circular dichroism, Δε M^-1^ cm^-1^, of unassembled (black line) and assembled strands (black line with red error bars) normalized with respect to total strand concentration. The red bars indicate the error in Δε found for several samples under different experimental conditions used to ensure complete assembly. C) Time courses of the assembly reaction monitored at 218 nm for total strand concentrations of (from top to bottom) 20, 40, 60, 80 and 100 µM. D) Free strand concentration as a function of time for samples at 30 µM total strand concentration at 10 °C (black), 20 °C (red) and 30 °C (blue).

Experiments were done at several high strand concentrations and prolonged incubations to produce fully assembled strands (Fig. 1B). These were normalized to molar circular dichroism, Δε = mdeg/(32980·C· l) M^-1^cm^-1^, with respect to strand concentration. For the assembled product, a value of -30.4±2.0 at 218 nm resulted. This value refers to the properties of a strand within the assembly. For the unassembled strands, Δε was determined for the spectrum of each strand separately, then taking their average, since by design all experiments presented in this study contained equimolar strand concentrations. A value of +10.9±0.7 at 218 nm was determined for the average unassembled strand molar circular dichroism. Analysis of these spectra by protein secondary structure analysis package BestSel ^21^ confirmed that the free and assembled spectra are consistent with those of unstructured peptide strands and a beta-sheet structure in the assembly. More specifically, BestSel indicates an assembly containing anti-parallel strands ^6^.

Panel C in Figure 1 shows primary data collected at 218 nm as a function of strand concentration. Full equilibration of these samples under experimental conditions was verified by careful examination of the slopes of the data over the last 10-20 minutes using first derivative analysis to ensure that it was zero within error. All samples were fully equilibrated by 240 minutes. From the observed Δε values at equilibrium the concentrations of unassembled and assembled strands are readily calculated as the linear combination of the limiting Δε values determined in Fig. 1B. This enables quantitative analysis of the thermodynamics and kinetics of the assembly reaction.

### Assembly reaction order

Figure S3 shows the results of an initial determination of the reaction order using the kinetic data of Fig. 1C. The analysis showed that over the 20-100 µM concentration range the disappearance of free strands was best fit by a sixth-order rate law, as judged by visual inspection of residual plots and by more sophisticated statistical tests as described in the figure caption. Elementary sixth-order reactions are thought to be impossible, since they require simultaneous collision of six reactants, but these data demand a sixth-order rate law to match the shape of the data curves. All attempts to fit lower reaction orders showed inferior fit statistics and systematic deviations of the residuals from a random Gaussian distribution.

The data of Figure 1C and D show that not only does assembly have the shape of a sixth-order reaction, but also that it is reversible. Samples at different concentrations and temperatures equilibrate to different extents of reaction and do not proceed to complete consumption of reactant strands. Such equilibration requires a finite reverse rate constant to balance the forward rate constant driving assembly.

Because reversible single-step sixth-order reactions are uncommon, simulated progress curves were used to develop intuition regarding the kinetic behavior expected for this effective mechanism (Fig. S4). Simulations employed assumed sixth-order forward and unimolecular reverse rate constants. The resulting time course has a complex appearance. Its initial portion displays a rapid burst phase of strand consumption, followed by a much slower apparent crawl toward equilibrium. For the simulated trace in Fig. S4A at an initial strand concentration of 60 µM, one-half of the total strands are consumed within 1.2 min. The first derivative of the simulated progress curve indicates that the slow phase is complete by 200–240 min. At that time, the system consisted of an equilibrium mixture of free strands and assembled hexamer, from which an equilibrium constant can be determined. Systematic variation of the forward rate constant at fixed reverse rate constant changes both the magnitude of the initial burst and the final extent of assembly (Fig. S4B). In contrast, variation of the reverse rate constant at fixed forward rate constant primarily changes the equilibrium endpoint (Fig. S4C). For comparison, Fig. S5 contrasts normalized simulated progress curves for reversible second- and sixth-order assembly reactions. The reactions were adjusted to have the same equilibrium extent of assembly and were plotted on a common dimensionless time scale. Relative to the familiar second-order dimerization reaction, the sixth-order process undergoes an exceptionally rapid initial rise followed by pronounced sensitivity to depletion of free strands. The inset expands the early portion of the curves and shows that the initial rise is continuous, but dramatically steeper for the sixth-order reaction. Detailed fitting of the experimental kinetic data to this mechanism is presented after the following sections on the characterization of the equilibrium thermodynamics of the assembly process.

### Thermodynamics of assembly

The minimal mechanism consistent with the data is for the assembly of complementary strands into a hexamer

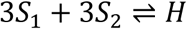

which, because we always maintain equal strand concentrations, reduces to

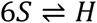

The equilibrium constant for the sixth-order assembly is

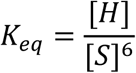

with units of *M*^−5^. The conservation equation for total strands is *S_T_* = *S_f_* + 6*H*.

Figure 2A shows the assembly isotherm for the association of octapeptides α-(M)_4_ and α-(C)_4_. The plot shows a highly cooperative assembly process with a characteristic “hockey stick” shape. There is a distinct lag before the onset of assembly at a critical assembly concentration near 7 µM total strand concentration. The cooperative self-assembly data were fit numerically using Origin software to a non-linear monomer–hexamer equilibrium isotherm derived from the Law of Mass Action, defined as H = *K_eq_*(*S_T_* − *nH*)*^n^*. The fit in Fig. 2A yielded (after scaling to 1M standard state) *K_eq_* = 2.3 ± 0.2 × 10^22^ *M*^−5^ and *n* = 6.1 ± 0.1. The best fit line and residuals are shown in the figure. The free energy, is ΔG^0^ = -RT*ln*K_eq_, =-30.5±0.1 kcal/mol.

**Figure 2.**
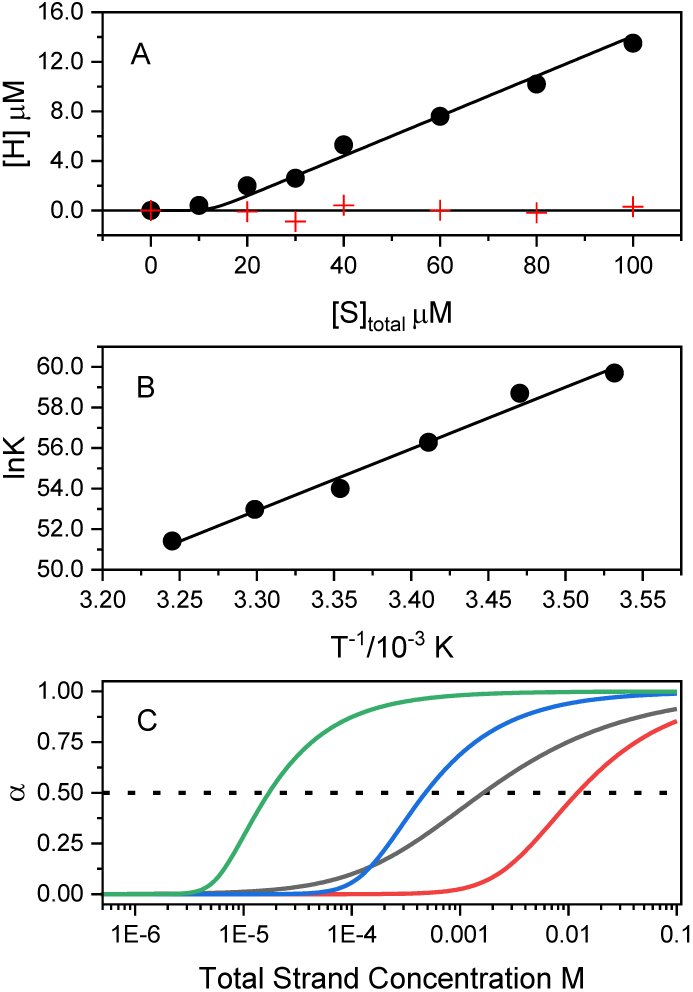
Equilibrium aspects of the assembly of octapeptides α-(M)_4_ and α-(C)_4_. A) Assembly of hexamer as a function of total strand concentration. The data (circles) fit the model described in the Supporting Information (red line) to yield the hockey stick shaped curve, with the residuals shown as the plus sign symbols around 0 on the x-axis. B) van’t Hoff plot for the assembly reaction. C) Comparison of the fraction of strands assembled (α) as a function of total strand concentration of representative duplex (blue), triplex (green), tetramolecular G-quadruplex (red) assemblies with the assembly of the octapeptides α-(M)_4_ and α-(C)_4_. The dotted line shows α = 0.5, the half-assembly point.

From the temperature dependence of a sample at 30 µM total strand concentration (Fig. 1D), K_eq_ at six temperatures over the range 10-35 °C equilibrium constants were determined and the van’t Hoff plot in figure 2B was constructed. The plot is linear and from the slope (-ΔH/R) an enthalpy of -60.4±4 kcal/mol was determined. The entropy, ΔS = (ΔH - ΔG^0^)/T, is - 100±13 cal/mol-K.

From this thermodynamic profile, the favorable free energy of the assembly reaction (-30.5 kcal/mol) is driven by the large negative enthalpy (-60.4 kcal/mol) and is opposed by an unfavorable entropy term TΔS (-29.8 kcal/mol). The unfavorable entropy reflects a battle between an enormous unfavorable conformational entropy and a favorable entropy contribution from the removal of solvent accessible surface area as per the hydrophobic effect. The conformational entropy arises from the loss of strand translational and rotational freedom (6 strands combining into one object) and from the restricted bond rotation in the PNA backbone strands within the assembly^22^. The resultant negative total entropy indicates the conformational entropy is incompletely balanced by the entropic contributions of surface area burial. The unfavorable total entropic cost of assembly is overcome by the large exothermic enthalpy. The favorable enthalpy likely arises from hydrogen-bond formation and stacking interactions among the mimetic bases in the rosette core, together with hydrogen bonding within the *β*-sheet structure of the strand backbones. Models of a cylindrical hexamer suggest that the core could be stabilized by up to 72 hydrogen bonds between paired bases, in addition to stacking interactions within the rosette core. The *β*-sheet backbone structure on the cylinder surface could contribute to tens of additional hydrogen bonds. These extensive hydrogen-bonding arrays can make a substantial favorable contribution to the enthalpy of hexamer formation, even after accounting for the exchange of solvent–solute interactions for internal hydrogen bonds. Best estimates for the net free energy of a hydrogen bond in DNA base pairs range from -0.7 to -1.6 kcal mol^-1^ ^23^. This energetic picture is fully consistent with a cooperative assembly process leading to hexamer formation.

The thermodynamic profile of hexamer assembly differs from that observed for the assembly of cyanuric acid (CA) and melamine (M), along with their water-soluble trivalent derivatives (TCA and TM) ^3^. The unfunctionalized parent heterocycles undergo spontaneous, unchecked noncovalent polymerization. This leads to the formation of macroscopically large, visible precipitates that quickly sediment out of the water solution. For the trivalent TCA/TM system with a flexible, charged synthetic scaffold, the assembly is more controlled and yields a uniform, stable nanoparticle suspension with low polydispersity, where the particles measure 100 to 200 nm in diameter. Both types of assembly are exothermic as measured by ITC, but complete thermodynamic profiles were not presented because a suitable equilibrium model could not be developed. The authors concluded, despite the lack of a full entropic profile, that assembly is driven by highly exothermic heteroaromatic ring-stacking coupled to an endothermic proton transfer, wherein polar surface burial overrides any classical hydrophobic effects. This mechanism was supported by solvent perturbation via Hofmeister salts and deuterium oxide (D_2_O) isotope effects.

Understanding the relative stabilities of multiple-stranded assemblies by comparison of their standard free energy changes is difficult and nonintuitive, since the magnitudes of the values vary significantly and it is difficult to understand how molecularity is factored in. Figure 2C shows an attempt to clarify the issue. The stabilities of duplex, triplex and quadruplex DNA were compared to the hexamer described here. To do so, nucleic acid assemblies containing base interactions of strands with 4 residues were assumed and arbitrarily chosen to contain only G or C bases. For tetramolecular G-quadruplex formation, experimental free energy values were available^10–11^ for the reaction 4*S* ⇌ *Q*. For duplex and triplex formation, free energies were estimated from established nearest-neighbor rules^24–27^. The fraction of strands assembled as a function of total strand concentration was estimated by converting the standard free energies for each type of assembly to its underlying equilibrium constant, then solving the mass action equilibrium to determine the distribution of free and assembled strands at each total strand concentration (see Methods in Supporting Information). Figure 2C shows the results, providing a more intuitive picture of stability. The plot shows the fraction of strands assembled at a given strand concentration for each. For these assemblies of comparable strand lengths, the hexamer described here is seen to be more stable than any of the others, with the tetrameric quadruplex following in stability. A subtlety is revealed that at low total strand concentration (up to 10 mM), duplex initially forms more readily than quadruplex, but after that point the greater cooperativity of quadruplex formation favors its assembly. Under the conditions of this comparison, the short strand lengths and the base composition are not favorable for triplex formation. The total strand concentration when half the strands are assembled, 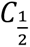, provides a convenient metric for ranking stability. 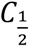 values show the ranking hexamer<quadruplex<duplex<triplex, with values of 17, 492, 1714, and 12500 µM, respectively. This exercise shows that the hexamer assembly is much more favorable than any other DNA assembly of similar strand length under similar solution conditions.

### Kinetics of assembly

Figure 3 shows an analysis of the kinetic data obtained at 60 µM total strand concentration for both the disappearance of strands and the formation of hexamer. The data show the expected burst and crawl phases expected for a reversible sixth-order reaction. The governing differential equations for the mechanism are

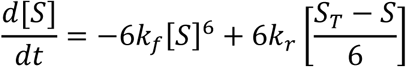

**Figure 3.**
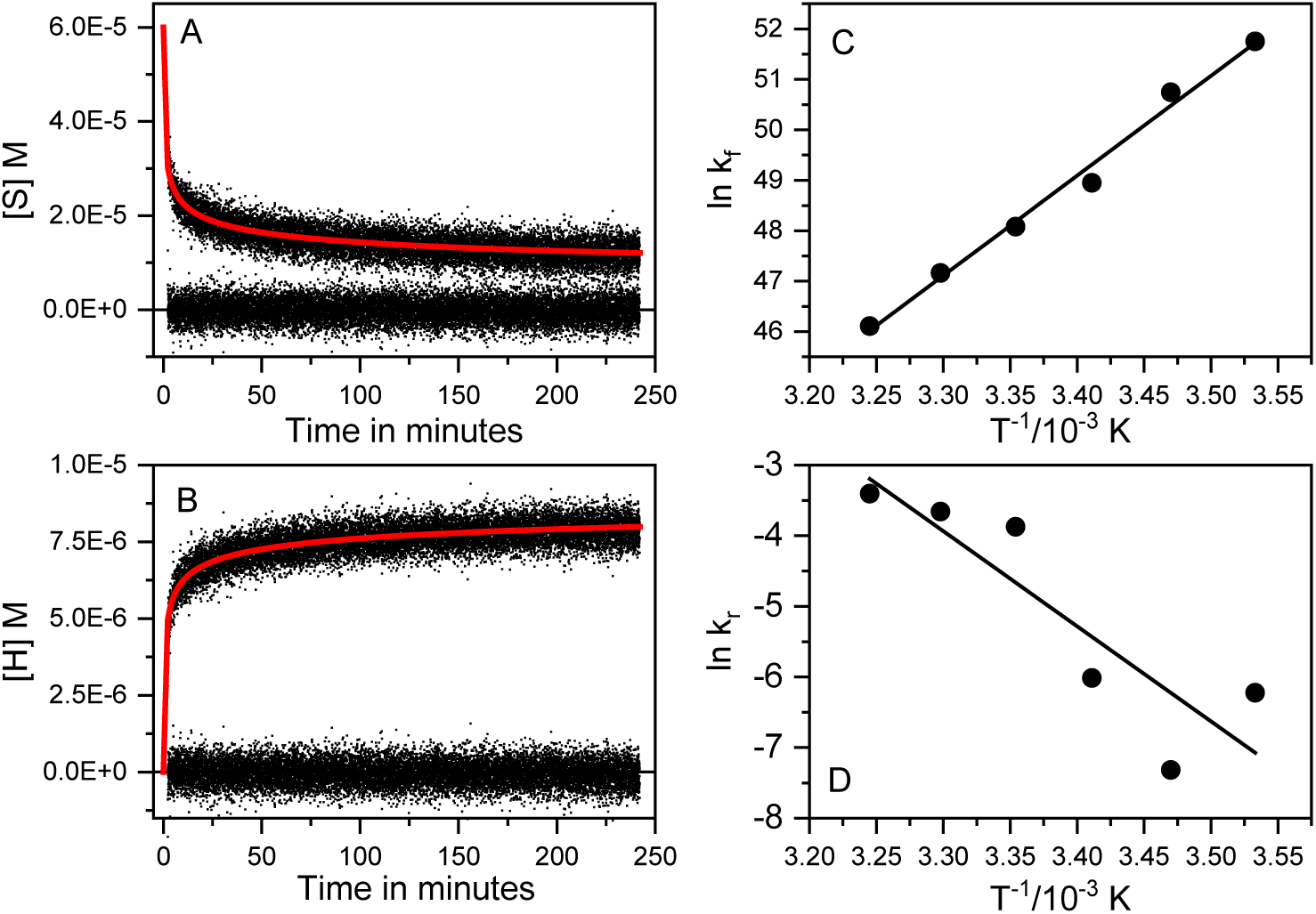
Kinetics of hexamer assembly. A) Disappearance of strands after mixing as a function of time at 25 °C and a total strand concentration of 60 µM. The red line is the fit to a reversible sixth-order reaction, with the residuals of the fit shown at the bottom of the panel. B) Same sample and conditions but showing the formation of hexamer as a function of time. C) Arrhenius plot for the forward sixth-order rate constant. The positive slope yields an unphysical activation energy E_a_ = -39.4±0.2 kcal/mol. D) Arrhenius plot for the reverse first-order disassembly rate constant. The negative slope yields an expected positive activation energy.

And

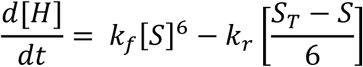

The fits of the data to these differential equations showed random residuals over the entire time course. From panel 3A values of k_f_=5.6±0.1×10^20^ M^-5^ min^-1^ and k_r_=1±0.6×10^-4^ min^-1^ were obtained for the disappearance of strands. From panel 3B values of k_f_=5.6±0.1×10^20^ M^-5^ min^-1^ and k_r_=0.2±0.1×10^-4^ min^-1^ were obtained for the appearance of hexamer. Clearly the forward rate constant estimates agree better than the reverse rate constants, due to enormous difference in magnitudes of the two rate constants leading to numerical stiffness in the fitting algorithm. All concentration-dependent kinetic traces from 20 to 100 µM are best described by a reversible sixth-order assembly model, but the nonlinear fits are ill-conditioned and yield only consistent order-of-magnitude estimates of the forward rate constant and very poorly constrained values of the reverse rate constant.

A more detailed exploration of the error space of parameter estimation is shown in Figure S6 that indicates that only an upper limit for k_r_ can be determined by the fitting algorithm. Further, there is a strong correlation between estimates of the forward and reverse rates. Nonetheless, this model provides an excellent fit to the shapes of all our data curves over the 20 – 100 µM concentration range and the 10-35 °C temperature range. It is extremely rare to observe primary data that follows sixth-order kinetics; indeed, we could find no other example after exhaustive search using the AI bibliographic search engine *Elicit™*, after screening and analysis of comprehensive kinetic compilations spanning multiple chemistry subdisciplines.

From the temperature dependence of k_f_ and k_r_ the Arrhenius plots shown in Fig. 3C,D were obtained. These plots reveal a complexity. While the plot for k_r_ (Fig. 3D) shows the expected negative slope corresponding to a positive activation energy, E_a_, the plot for k_f_ has a positive slope, corresponding to a *negative* activation energy of -39.4±0.2 kcal/mol. This is inconsistent with the behavior of an elementary reaction step and indicates hidden complexity in the assembly process, with either faster preequilibrium steps, steps undetectable by CD, or both. While such behavior is unusual, it is not unprecedented and was observed in the assembly of multimeric strands into a parallel G-quadruplex structure^10–11^. While the observed progress curves are fully consistent with a macroscopic reversible sixth-order assembly, the negative E_a_ signals that a more complex reaction mechanism is needed to fully account for the data.

### Relaxation kinetic analysis

Before exploring more complex mechanisms, we examined the robustness of the reversible sixth-order fit by an independent analysis based on relaxation kinetics. The results are shown in Figure S7. The relaxation time for the approach to equilibrium was determined by fitting each time course to biexponential decay and assigning the slower decay to the approach to equilibrium relaxation time, *τ*. This enables linearization of the full differential equation for an *n*-th order reversible assembly reaction approaching equilibrium, yielding the equation

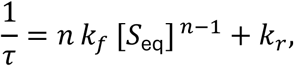

where [*S*_eq_] is the concentration of free strands at equilibrium^28^. A plot of 1/*τ* versus [*S*_eq_]*^n^*^−1^ for *n* = 6 (Fig. S7) is linear and gives from the slope *k_f_* = (8.6 ± 0.2) × 10^19^ M^−5^ min^−1^ and from the intercept *k_r_* = 0.01 ± 0.002 min^−1^, which are consistent in order of magnitude with the values obtained from full ODE fitting but are much more precisely defined. In contrast to the individual concentration nonlinear ODE fits, which yield strongly correlated and ill-conditioned estimates of *k_f_*and *k_r_*, the relaxation analysis is based on a single global linear relation between the observed relaxation rates and concentration, so it provides much more stable and statistically reliable values for the rate constants. Fits of the data to the relaxation equation with *n* < 6 were systematically statistically inferior. This alternate approach supports the conclusion that the effective reaction order is six.

Notably, the relaxation analysis yields an equilibrium constant 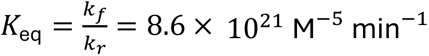 10^21^M^−5^ min^−1^ that agrees, within a factor of 2-3, with the value determined independently from the equilibrium “hockey-stick” isotherm (Fig. 2A).

Taken together, these results indicate that the observed sixth-order kinetics provide a robust macroscopic description of the assembly dynamics, but that the underlying microscopic mechanism must contain one or more additional step because of the abnormal Arrhenius behavior. We therefore considered more detailed kinetic schemes, beginning with a minimal pre-isomerization mechanism, followed by a commonly assumed sequential oligomerization model that is often invoked for cooperative, multimeric assembly processes.

### Isomerization of strands preceding assembly

A “preisomerization” model provides the mosteconomical mechanism that remains consistent with the two-state CD observables and the cooperative equilibrium isotherm, while introducing only the minimal number of additional steps needed to restore a physically reasonable activation barrier for assembly. This simple extension assumes that a rapid conformational isomerization of the strands precedes hexamer formation:

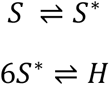

The first step has rate constants k_1_ and k_-1_ for the forward and reverse steps; the second assembly step has k_f_ and k_r_ for the assembly process.

Figure 4 shows the results of fits to this model. From fits of the data over the concentration range 20-100 µM, the values of k_1_ = 0.65±0.2 min^-1^and k_-1_=0.9±0.25 min^-1^ were obtained for the first-order preisomerization step. From these the derived quantities K_1_= k_1_ / k_-1_=0.7±0.2 and a relaxation time of 1/τ= k_1_ + k_-1_=1.6±0.4 min were obtained. From the equilibrium constant K_1_, a free energy change of +0.2 kcal mol^-1^ is calculated, indicating that the strand isomerization is slightly unfavorable. Figure 4B shows a van’t Hoff plot for the isomerization reaction over the 10-35 °C range, yielding an enthalpy of -12.3 kcal mol^-1^, indicating that increased temperature could drive the isomerization step.

**Figure 4.**
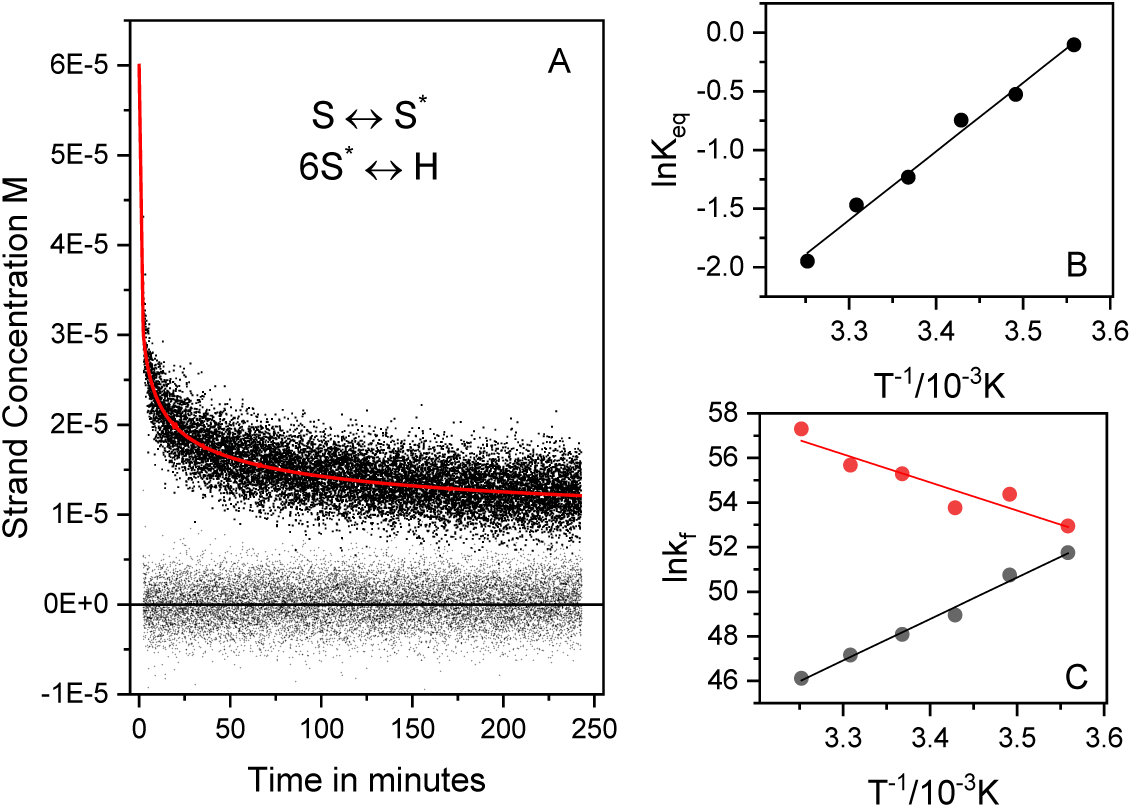
An alternate kinetic model. A) Fits of the same data in figure 3 to a two-step kinetic mechanism in which a fast stand isomerization is followed by a reversible sixth-order assembly reaction to form a hexamer. B) van’t Hoff plot of the temperature dependence of the stand isomerization equilibrium constant. The positive slope indicates an exothermic isomerization with negative enthalpy. C) Contrasting Arrhenius behaviors of the forward assembly rate constants for this preisomerization model (red symbols) and the single-step reversible sixth-order assembly. The preisomerization model yields a physically reasonable activation energy of +26.6 kcal/mol for the forward assembly rate constant.

Fits to the model yield values of k_f_ and k_r_ for the second sixth-order assembly step that are consistent with the magnitudes of values obtained from the single-step mechanism shown in Fig 3. But, as seen in Fig. 4C, rate constants for the assembly step now show the expected Arrhenius behavior, yielding a positive activation energy for k_f_ of +26.6 kcal/mol.

While the preisomerization analysis is numerically challenging, it nevertheless provides important qualitative insights into the assembly mechanism. Fits to the scheme describe the data as well as the simpler single-step model, but the additional microscopic rate constants are not tightly constrained. The preisomerization rates show substantial variation between experiments and conditions, and formal statistical criteria comparing the two models suggest that the extra parameters are at best only marginally warranted. In that sense, the current data does not allow precise microscopic parametrization of the preisomerization step or the subsequent assembly.

Even with these limitations, the preisomerization model delivers physically meaningful thermodynamic signatures that strongly suggest the presence of added complexity in the assembly landscape, with faster steps than we cannot measure by out method or with intermediates undetectable by CD. The temperature dependence of the assembly rate yields a positive activation energy of the expected magnitude for the assembly, and the equilibrium constant for the preisomerization step displays a clear, enthalpically driven temperature dependence. Together, these features indicate the presence of a genuine preassembly process that precedes the high-order growth step, even if its individual microscopic rate constants are only loosely determined.

### A sequential assembly mechanism can manifest as a macroscopic sixth-order progress curve

Cooperative multimeric assembly reactions are commonly explained by a stepwise oligomerization scheme in which monomers add sequentially to growing intermediates, with each step governed by its own forward and reverse rate constants^29–30^. The mechanism is appealing because it avoids the physical implausibility of a single-step elementary sixth-order reaction, which would demand the simultaneous collision of six independent reactants. A recent example used single-molecule methods to unravel the mechanism of hexameric insulin assembly using a similar sequential framework^31^. However, as that example highlights, even a minimal sequential model for hexamer assembly quickly becomes unwieldy, requiring 10 or more microscopic rate constants. For the hexameric assembly studied here, the information content of macroscopically observed progress curves is insufficient to uniquely determine or reliably fit that many parameters. Direct unconstrained fitting of such a model would inevitably lead to severe parameter correlations and non-unique solutions, providing little structural insight beyond what is already captured by the simpler effective sixth-order law. Furthermore, while intermediate oligomeric species in a sequential model might be expected to possess distinct spectroscopic signatures, our singular value decomposition (SVD) analysis of the circular dichroism (CD) spectra reveals strictly two-state behavior.

Despite these limitations, kinetic simulations can provide crucial insight into whether a highly constrained sequential mechanism can successfully manifest as a macroscopic sixth-order progress curve while remaining consistent with our two-state spectroscopic data. The sequential hexamer assembly model is formulated as follows:

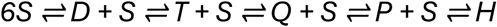

where S, D, T, Ǫ, P, and H represent the monomeric strand, dimer, trimer, tetramer, pentamer, and hexamer, respectively. This mechanism is governed by 10 independent parameters comprising the forward (kf,i) and reverse (kr,i) rate constants for each step i. To test the feasibility of this pathway without overparameterization, we implemented a heavily constrained non-linear least-squares optimization in MATLAB (see Supporting Information for implementation details). To regularize the fit and drastically reduce the degrees of freedom, the intermediate breakdown rates were fixed to a single, highly unstable common value (kr,1 to kr,4 = 100 min⁻¹). These are reasonable values given that the dissociation rate constants of DNA, RNA and DNA-RNA hybrid oligonucleotides ranged from 0.5 to 500 s^-1^, with forward bimolecular association constants clustered more tightly around 10^6^ M^-1^s^-1^ ^32^.

Figure 5 demonstrates the success of this constrained optimization. Data are shown for the same 60 µM sample shown in previous figures. As shown in Fig. 5A and B, the simulated trajectories for both the monomeric strand (S) and the final hexamer (H) overlay with the experimental progress curves, capturing the characteristic apparent sixth-order decay profile. Crucially, the corresponding species plots (Fig. 5C,D) reveals that all transient intermediates (D, T, Ǫ, and P) remain at low, sub-micromolar steady-state concentrations throughout the entire time course. Because these intermediates are never significantly populated, they would remain undetected by our CD method (even if they had distinctive spectra), reconciling the sequential mechanism with the two-state behavior observed in our SVD analysis. The downstream apparent macroscopic forward and reverse rate constants from the optimized microscopic parameters can be calculated from the equations yields.

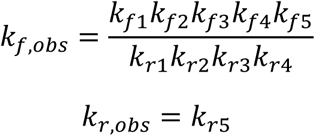

**Figure 5.**
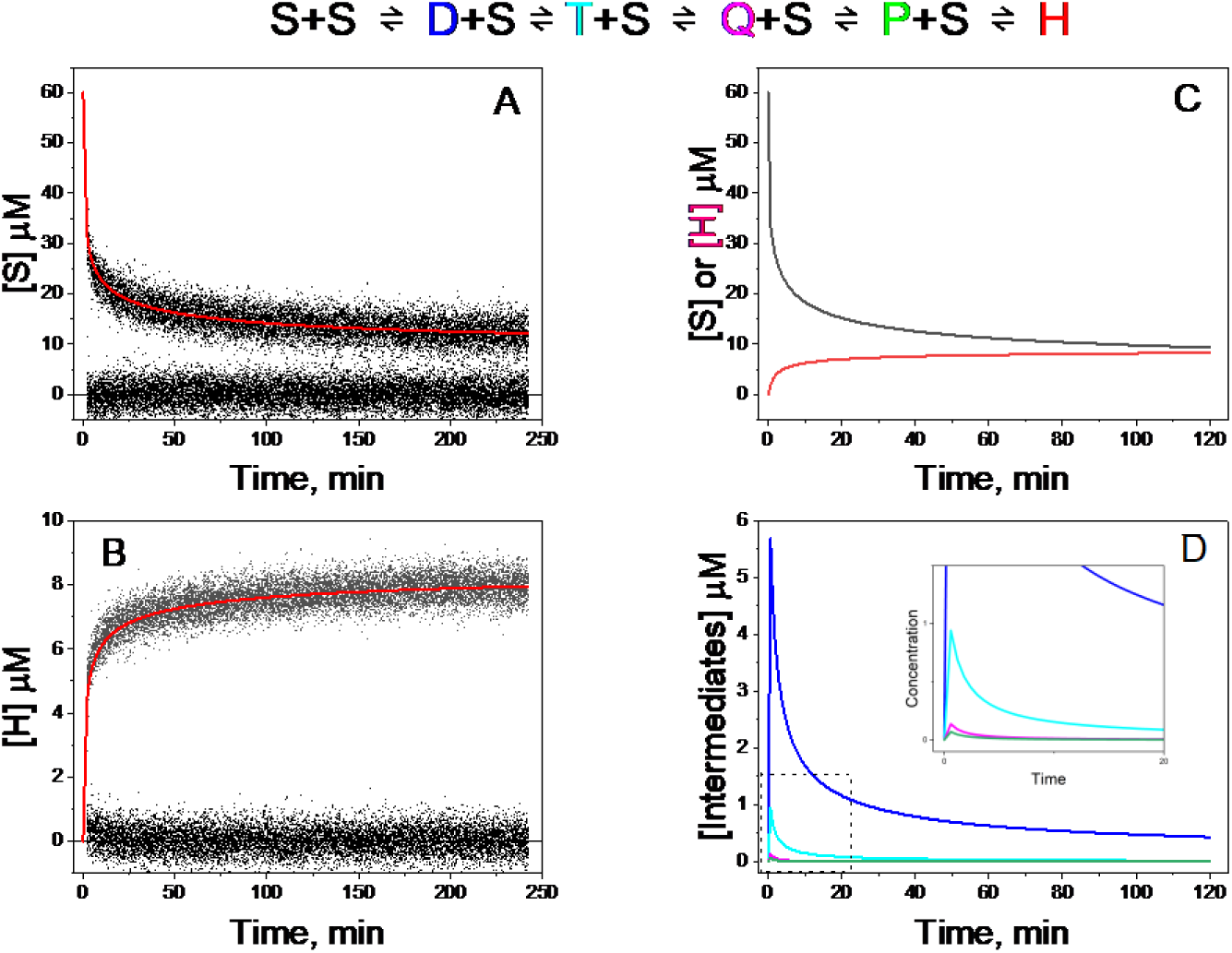
Results of simulations of sequential assembly model. A) Computed model (red line) for the disappearance of strands (60 µM) with time. The residuals are shown on the line at 0. B) Computed model (red line) for the formation of hexamer. B, C) Species plots showing the changes in concentration with time for strands and hexamer (C) and intermediates (D). The curves in B are the “fit” curves in panels A and B. The intermediate curves in (D) are color-coded with respect to the reaction model above. The inset in (D) expands the first 20 minutes for clarity. These simulation results show that the sequential model can collapse into an apparent two-state, reversible, sixth-order progress curve (A, B) with no detectable intermediates for particular sets of microscopic sequential rate constants. The general requirements for this are a slow initial dimerization step, unstable intermediates in low, near steady-state concentrations and a final bottle neck with a rapidly formed, but slowly dissociating, hexamer. The exact rate constants used for this simulation are shown in Table S1.

Yielding values of 7.0×10^20^ M⁻⁵min⁻¹ and 1.2×10^-4^ min^-1^. These values compare to the fitted parameters from the simpler two-state model of 5.6×10^20^ M⁻⁵min⁻¹ and 0.2×10^-4^ min^-1^ (Fig. 3). The optimized values for the microscopic rate constants are shown in Table S1. Simulations were also done using data over the range of 20 -100 µM with excellent agreement. Additonal simulations using a range of 5-500 min^-1^ for the fixed dissociation rate constants of the intermediates also produced apparent sixth-order macroscopic progress curves, with the lower limit just beginning to show systematic deviations between the experimental and simulated progress curves.

Given the constraints of the available macroscopic data, this set of microscopic parameters represents only one possible set of values rather than a definitive unique solution. Nonetheless, these simulated results serve as a powerful proof-of-concept. They demonstrate that a physically plausible, multi-step pathway under realistic limiting conditions can collapse to reproduce a rare macroscopic sixth-order progress curve without requiring an implausible, single-step, elementary collision.

### The precedent of tetrameric G-quadruplex assembly

The assembly of oligomeric G strands into a four-stranded parallel G-quadruplex structures^8, 10–12^ provides another illustration of sequential assembly masking as a simpler higher-order progress curve. Those studies show slow assembly curves that were fit to fourth-order rate laws and forward rate constants that displayed negative activation energies. Rate constants were highly dependent on DNA versus RNA, temperature, concentration, ionic strength, and strand length. Collectively, a vast range in forward rate constants was reported, from 6.1×10^4^ to 5×10^12^ M^-3^s^-3^. In addition to these studies, temperature-jump kinetic studies were done to measure the rate constant for disassembly of tetrameric G-quadruplexes ^33^. Estimates of the reverse rate constant in that study varied in magnitude from 10^-10^ s^-1^ at physiological temperatures to 10^-4^ s^-1^ at elevated temperatures. Freire^11^ recognized that the unphysical Arrhenius behavior masked hidden complexity, and proposed a two-step reaction mechanism with strand dimerization followed by bimolecular assembly of the dimer intermediate into the quadruplex. No attempt was made to quantitatively analyze experimental data to extract microscopic rate constants. Hardin^8^ described an Koshland-Nemethy-Filmer sequential, induced-fit, allosteric model for the assembly process. An especially careful NMR study by Bardin and Leroy found that a sequential assembly mechanism, with a rate-limiting triplex formation step, provided the best description of tetramolecular G4 assembly^12^. The overall reaction remained fourth-order. These studies show that now there are at least two cases, ours and tetramolecular G4 formation, that contradict the kinetic dogma that “…equations of the fourth order and higher orders do not even need to be considered”.

### Contrasting behavior of insulin hexamer assembly

The recent single-particle analysis of insulin hexamerization illustrates the type of experiment required to resolve a full microscopic assembly network ^31^. Direct fluorescence measurements identified monomers through hexamer states and numerous individual association and dissociation transitions that are necessarily averaged in bulk observables. In contrast, the present CD/SVD measurements establish a well-defined two-state macroscopic description and place strong constraints on possible mechanisms, but cannot identify low-population intermediates or resolve events faster than the experimental time resolution. Definitive discrimination among alternative hidden pathways will require new observables and greater temporal resolution, ideally in experiments that can directly detect individual oligomeric states.

### Conclusions

Structure determinations provide a static picture of the [α-(M)_4_/α-(C)_4_]_3_ hexamer, while these thermodynamic and kinetic studies explore why and how hexamer forms. Circular dichroism and singular-value decomposition show that complementary α-(M)_4_ and α-(C)_4_ strands assemble into a highly stable, enthalpically driven hexamer with two-state behavior and no detectable intermediates. Kinetic studies of hexamer formation establish that its assembly are accurately described by a reversible sixth-order macroscopic rate law. A negative apparent activation energy of the forward rate constant is inconsistent with an elementary six-strand collision, pointing to hidden complexity. Analysis with a preisomerization model and simulations of a constrained sequential model demonstrate how hidden conformational or oligomerization steps, including low-population and spectroscopically silent intermediates, can collapse into an apparent two-state, sixth-order progress curve. This system provides an unique example of a highly cooperative hexamer formation can manifest as a deceptively simple sixth-order macroscopic progress curve, concealing a more complex underlying microscopic reaction network.

The thermodynamic profile of hexamer assembly shows that the reaction is driven by a large negative enthalpy and is opposed by entropy. That pattern is consistent with formation being driven by favorable molecular interactions (H-bonding, stacking) along with hydrophobic forces from the burial of solvent accessible surface area. The opposing entropy is dominated by the cost of the loss of rotational and translational freedom that results from combing free single strands into a constrained hexamer. An additional large entropic cost comes from the immobilization of bonds in the α-DPNA backbone. PNA is known to be more flexible than DNA and RNA, hampering its binding selectivity^34^. This raises a design consideration when attempting to tether interactive mimetic base to backbones in hopes of constructing complementary polymers. Complementary strand interactions always come with a significant energetic cost from the immobilization bond rotation of otherwise flexible backbones. For example, Searle and Williams decomposed helix-formation free energies into approximate physical contributions and estimated an adverse free-energy cost, primarily entropic, of 1.9–5.4 kJ mol⁻¹ (1 kJ = 0.24 kcal) per restricted phosphate-backbone rotor ^22^. Design of related polymersthat includes strand preorganization^34–35^ (although a daunting task) could further optimize hexamer assemblies for materials applications and provide insights regarding possible prebiotic polymers that were even more favored to self-assemble prior to the emergence of RNA.

## Acknowledgments

This work was supported by The Alfred P. Sloan Foundation (grant no. G-2021-16757) and by NASA Exobiology (19-EXO19-0018).

## AI Declaration

The authors used Perplexity AI (Perplexity AI, Inc.) as an interactive computational and editorial aid to assist with mathematical formulation and checking of analyses, statistical-analysis planning, development and debugging of computational scripts, and refinement of manuscript text. All scientific decisions, data processing, calculations, code implementation, interpretation of results, verification of cited literature, and final writing and editing were performed and independently reviewed by the authors. The authors take full responsibility for the accuracy, originality, and integrity of all content in this manuscript.

