## Supporting Information for "A unique sixth-order reversible assembly of hexameric mimetic nucleic acid"

### A macroscopic reversible sixth-order assembly of α-DPNA strands with nucleobases into a hexamer

**<u>CONTENTS</u>**

I. Materials

II. Methods

III. Figures and Tables

Figure S1. *Results of SVD analysis*

Figure S2. *Data reconstruction from SVD*

Figure S3. *Determination of reaction order*

Figure S4. *Simulations of a reversible sixth-order reaction*

Figure S5. Normalized kinetic progress curves for reversible dimer and hexamer assembly

Figure S6. *Mapping the error space of $k_r$*

Figure S7. *Relaxation kinetic analysis of the reversible sixth-order assembly reaction*

Table S1. Optimized microscopic rate constants for sequential assembly model

### I. Materials

Detailed synthesis, purification, characterization and stock solution preparation protocols of the α-DPNAs, α-(M)$_4$ and α-(C)$_4$ were previously reported. (in review). Sodium phosphate was prepared by mixing sodium phosphate monobasic monohydrate ($NaH_2PO_4 \cdot H_2O$) and sodium phosphate dibasic heptahydrate ($Na_2HPO_4 \cdot 7H_2O$) purchased from BDH. NaOH and $NH_4OH$ were used to adjust the pH of buffer and sample.

### II. Methods

#### Concentration Determination

Stock solutions of α-(M)$_4$ and α-(C)$_4$ were prepared by HPLC purification, followed by lyophilization and resuspension in $D_2O$ with the minimal amount of $NH_4OH$ needed for dissolving each oligomer (which were initially in a less-soluble acidic state due to the acidity of the HPLC running buffer). Oligomer concentrations were determined by the addition of a known amount of isopropyl alcohol (IPA) that was used as an internal concentration standard for analysis by $^1$H NMR. IPA was then removed by lyophilization and resuspended the oligomers in a precise volume of $H_2O$.

#### *Mixing Protocols*

Two sample preparation procedures were used. **Method 1 (high concentration)** was designed to maximize assembly: 1 µL aliquots of 10 mM α-(M)$_4$ and α−(C)$_4$ were added to 40 µL sodium phosphate buffer (pH 7), vortexed, diluted with water to a final volume of 200 µL, and vortexed again. This protocol produced final concentrations of 50 µM for each strand in 200 mM sodium phosphate buffer at a final pH of 7.2. Thus, α-(M)$_4$ and α−(C)$_4$ assembled at higher strand concentration were each initially 0.24 mM before dilution. **Method 2 (low concentration)** was used for kinetic and equilibrium measurements. Separate solutions of each strand were prepared at twice the desired final concentration in sodium phosphate buffer, then mixed in equal volumes to obtain the desired final strand concentrations and a constant final sodium phosphate concentration of 0.2 M. For example, equal 100 µL aliquots of 100 µM α-(M)$_4$ and α−(C)$_4$ yielded a 200 µL sample that was 50 µM in each strand. In contrast to Method 1, in this low concentration method strand assembly began only after the two strands were combined at their final, lower concentration.

Time- and concentration-dependent single-wavelength and full-spectrum CD measurements with real-time data acquisition used Method 2. Samples were mixed at room temperature, vortexed, immediately transferred to the CD cuvette, and data was recorded without delay. Mixing, cuvette filling, and initiation of data collection required approximately 2 minutes, which defined the dead time of the kinetic experiments.

For temperature-dependent kinetic measurements, the strands were mixed and transferred to a cuvette that was pre-equilibrated at the specified temperature for 10 minutes.

#### *CD Measurements*

All CD measurements were performed using a Jasco J-810 spectropolarimeter equipped with a Peltier temperature-controlled cell holder. Deionized water (18.2 MΩ/cm) from a Barnstead NANOpure Diamond water-purification system was used for sample preparation and experimental studies. Quartz cuvettes with 1 mm and 1 cm pathlengths were used to record CD spectra.

For conventional CD measurements, spectra were collected from 200 to 300 nm at a scan rate of 100 nm/min, a response time of 1 sec, data pitch of 1 nm, and a bandwidth of 4 nm. Four scans were accumulated and averaged, and buffer blank baseline subtracted. CD spectra were converted to molar circular dichroism, $\Delta\varepsilon$, using

$$\Delta\varepsilon = \frac{\theta}{32980cl}$$

where $\theta$ the CD signal in millidegrees, $c$ the molar concentration of strands, and $l$ is the cuvette pathlength in cm.

#### *Data Analysis and Fitting*

Nonlinear least-squares fitting and statistical analyses were accomplished using OriginPro, Version 2022 (OriginLab Corporation, Northampton, MA, USA). Data were fit to differential equations using the Origin App "Fit ODE" (v. 1.4) using a Runga-Kutta integration algorithm. Kinetic simulations were accomplished using MATLAB® version R2013b (The MathWorks Inc., Natick, Massachusetts, https://www.mathworks.com). The script used for simulation of the sequential model is provided as an *.m file in Supporting Information.

#### *Equilibrium Assembly Calculations*

Each nucleic-acid structure was represented as a concerted n-strand association,

$$\mathrm{n}S \rightleftharpoons S_\mathrm{n},$$

where n = 2, 3, 4, and 6 for the duplex, triplex, quadruplex, and hexaplex, respectively. At 298.15 K, the standard equilibrium constant was calculated from the assigned standard Gibbs energy according to

$$K_n^\circ = \exp[-\Delta G_n^\circ/(RT)],$$

where R = 1.9872 × $10^{-3}$ kcal $mol^{-1}$ $K^{-1}$. Assuming ideal dilute-solution behavior and a standard concentration c° = 1 M, the thermodynamic equilibrium constant was expressed as

$$K_n^\circ = ([S_n]/c^\circ)/([S]/c^\circ)^n,$$

and the corresponding concentration-form association constant was

$$Kc_{,n} = K_n^\circ/(c^\circ)^{n-1},$$

$$[S_n] = Kc_{,n}[S]^n.$$

When all concentrations were entered numerically in molar units, $Kc_{,n}$ had units of $M^{n-1}$ but the same numerical value as $K_n°$. For each total strand concentration, $C_t$, the equilibrium free-strand concentration, s = [S], was obtained from the strand mass balance

$$C_t = s + n[S_n] = s + nKc_{,n}s^n.$$

The unique physically meaningful root in the interval $0 \le s \le C_t$ was found numerically from

$$nKc_{,n}s^n + s - C_t = 0.$$

The fraction of total strand units incorporated into assembled structures was calculated as

$$f_{assembled} = n[S_n]/C_t = nKc_{,n}s^n/C_t = 1 - s/C_t.$$

The standard free energies and resulting equilibrium constants used in the calculations were as follows:

| **Assembly** | **n** | $\Delta G^{\circ}_{25C}$ **(kcal mol$^{-1}$)** | $Kc_{,n}°$ **at 298.15 K** | $Kc_{,n}$ **units** |
|---|---|---|---|---|
| Duplex | 2 | −3.5 | $6.10 \times 10^2$ | $M^{-1}$ |
| Triplex | 3 | −4.6 | $9.08 \times 10^3$ | $M^{-2}$ |
| Quadruplex | 4 | −14.0 | $1.83 \times 10^{10}$ | $M^{-3}$ |
| Hexamer | 6 | −33.5 | $3.59 \times 10^{24}$ | $M^{-5}$ |

.

Values of $\Delta G°_{25°C}$ for the hexamer were measured here and taken from the literature for the quadruplex. For the duplex a sequence of 5'GGGG-CCCC was assumed and the free energy calculated by nearest-neighbor rules. For the triplex formation the free energy of addition the third strand CCCC was calculated by triplex nearest-neighbor rules and the free energy of the overall reaction $3S \leftrightharpoons T$ was obtained as the sum of free energies of the two-step process

$$S_1 + S_2 \rightleftharpoons D \quad \Delta G_1$$

$$D + S_3 \rightleftharpoons T \quad \Delta G_2$$

$$3S \rightleftharpoons T$$

$$\Delta G_T = \Delta G_1 + G_2$$

#### *Kinetic Modeling and Mass-Action Differential Equations*

To describe the assembly kinetics of the α-dipeptide nucleic acid (αDPNA) system into hexameric structures, a stepwise mass-action kinetic model was developed. The assembly process is modeled via consecutive addition of monomeric strands ($S$) through transient intermediate states: dimer ($D$), trimer ($T$), tetramer ($Q$), and pentamer ($P$), culminating in the formation of the final hexamer product ($H$).

The system is governed by the elementary reaction network:

$$S + S \leftrightarrow D \quad \text{(forward k\_f1, reverse k\_r1)}$$

$$D + S \leftrightarrow T \quad \text{(forward k\_f2, reverse k\_r2)}$$

$$T + S \leftrightarrow Q \quad \text{(forward k\_f3, reverse k\_r3)}$$

$$Q + S \leftrightarrow P \quad \text{(forward k\_f4, reverse k\_r4)}$$

$$P + S \leftrightarrow H \quad \text{(forward k\_f5, reverse k\_r5)}$$

Applying the law of mass action yields a system of six coupled ordinary differential equations (ODEs) tracking the time-dependent concentration trajectories (μM) of each individual species over time:

$$d[S]/dt = -2r1 - r2 - r3 - r4 - r5$$

$$d[D]/dt = r1 - r2$$

$$d[T]/dt = r2 - r3$$

$$d[Q]/dt = r3 - r4$$

$$d[P]/dt = r4 - r5$$

$$d[H]/dt = r5$$

Where the individual reaction step velocities (r1 through r5) are explicitly defined by the difference between the forward and reverse elementary pathways:

$$r1 = k\_f1 * [S]^2 - k\_r1 * [D]$$

$$r2 = k\_f2 * [D] * [S] - k\_r2 * [T]$$

$$r3 = k\_f3 * [T] * [S] - k\_r3 * [Q]$$

$$r4 = k\_f4 * [Q] * [S] - k\_r4 * [P]$$

$$r5 = k\_f5 * [P] * [S] - k\_r5 * [H]$$

The model includes the following assumptions:

1. *Symmetric Intermediate Dissociation*: The breakdown rates of all transient pre-nucleation intermediates (*D*, *T*, *Q*, *P*) back to their constituent components are assumed to be identical and independent of oligomeric state. This structural simplification reduces parameter dimensionality and ensures numerical stability during convergence loops.
2. *Reversible Final Sink*: The final hexamer step is treated as a highly stable, independent reversible sink, governed by its own independent reverse rate constant (k_r5).
3. *Conservation of Mass*: No secondary aggregation pathways, off-target misfolding, or multi-body collisions occur outside of consecutive monomer addition.
4. *Ideal Solution Conditions*: Activity coefficients are taken as unity, allowing chemical activities to be represented directly by molar concentrations.

The MATLAB® script used to implement the model is provided in the Supporting Information as the file: *Sequential_Model.m*.

Specific details of the script include:

1. *ODE Integration*: The stiff differential equation network was simulated across time domains using the variable-order, numerical differentiation formula solver ode15s. Relative and absolute error integration tolerances were bounded at $10^{-6}$ and $10^{-8}$, respectively, to prevent numerical drift.
2. *Simultaneous Global Parameter Estimation*: A 6-parameter vector (kf1, kf2, kf3, kf4, kf5, and kr5) was optimized via a simultaneous global curve fit utilizing the trust-region-reflective nonlinear least-squares algorithm via lsqcurvefit. Rather than fitting individual trajectories independently, a single unified set of rate constants was forced to satisfy the coupled system across all time domains. The objective function minimized the global residual sum of squares across the shared experimental and simulated curves for both the monomer ([$S$]) and hexamer ([$H$]) channels simultaneously:

$$\min_p \sum \left[ \left([S]_{exp,i} - [S]_{mod,i}(p)\right)^2 + \left([H]_{exp,i} - [H]_{mod,i}(p)\right)^2 \right]$$

3. *Parameter Scaling & Bounds*: Search limits were restricted between a lower bound of $10^{-3}$ μ$M^{-1}$ $min^{-1}$ (0 for k_r5) and an upper bound of 50.0 μ$M^{-1}$ $min^{-1}$ (1.0 $min^{-1}$ for k_r5) to prevent unphysical negative values or runaway scaling.

#### *Macroscopic Kinetic Rate Translation*

To compare the microscopic parameter values against conventional high-order reaction mechanisms, an apparent macroscopic forward assembly rate constant (k_f_macro) was computed from the steady-state values of the microscopic parameters under rapid intermediate equilibrium:

$$k\_f\_macro = (k\_f1 * k\_f2 * k\_f3 * k\_f4 * k\_f5) / (k\_r\_frozen)^4$$

The resulting k_f_macro values were converted from working units (μ$M^{-5}$ $min^{-1}$) into absolute standard SI units ($M^{-5}$ $min^{-1}$) for direct benchmark verification against fits to the simple two-state models described in the text.

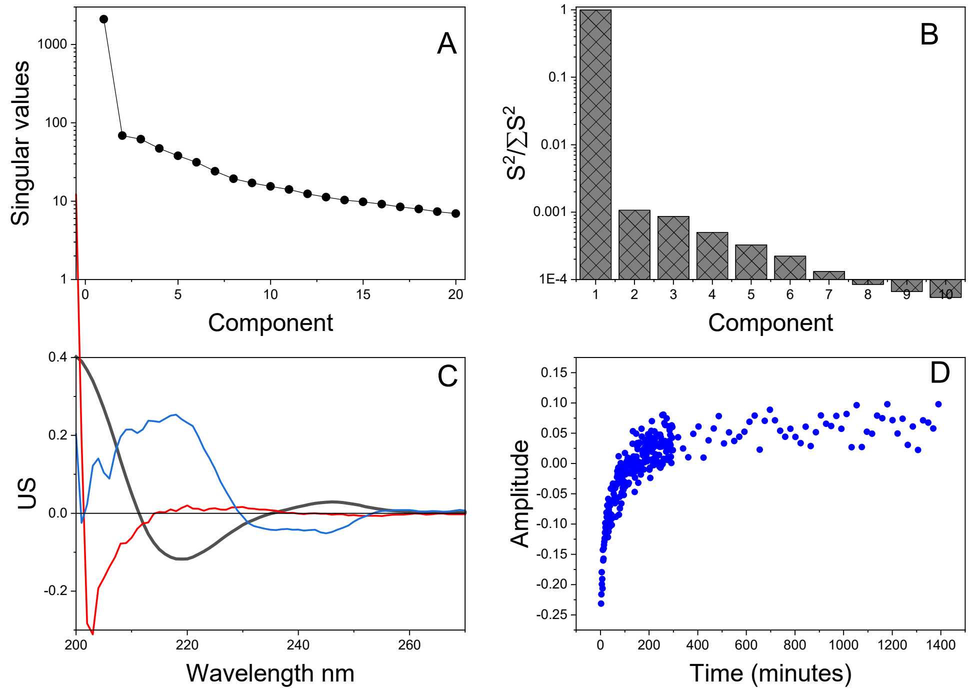


**Figure S1**. *Results of SVD analysis*. Singular value decomposition (SVD) was used to analyze the complexity of the primary kinetic data shown in Figure 1 using published methods[1-4]. The primary data were recast as a 101 × 294 matrix **D**, with rows as wavelength and columns as time. SVD factorizes the data matrix into three matrices, $\mathbf{D=SUV^T}$. **S** is a diagonal matrix containing the singular values ordered from largest to smallest, which represents the importance of each basis vector. **U** contains the basis spectra of the family of spectra in the primary data. **V** contains the amplitude vectors that weigh each component. Panel A shows the first 20 singular values, presented as a semilog plot for clarity. Panel B is a Scree plot to quantitate the contribution of each singular value (for the first 10 components); again, note the semilog plot for clarity. These results show that 3 components account for 99.8% of the variance in the primary data. The spectral shapes of these components are shown in panel C. The primary component is of the black curve, the second component the red curve and the third component the blue curve. The black and red curves are of the final and initial spectral shapes, respectively. The blue curve is the difference between these two and maps the shape of the spectral change over time in the kinetic experiment. Panel D show the amplitude of this change over time, representing a kinetic time course for the reaction over the entire spectral wavelength range. These results show the primary data contain only two significant spectral species, and their difference. This means that the reaction is a simple two-state process, with no detectable intermediate spectral species.

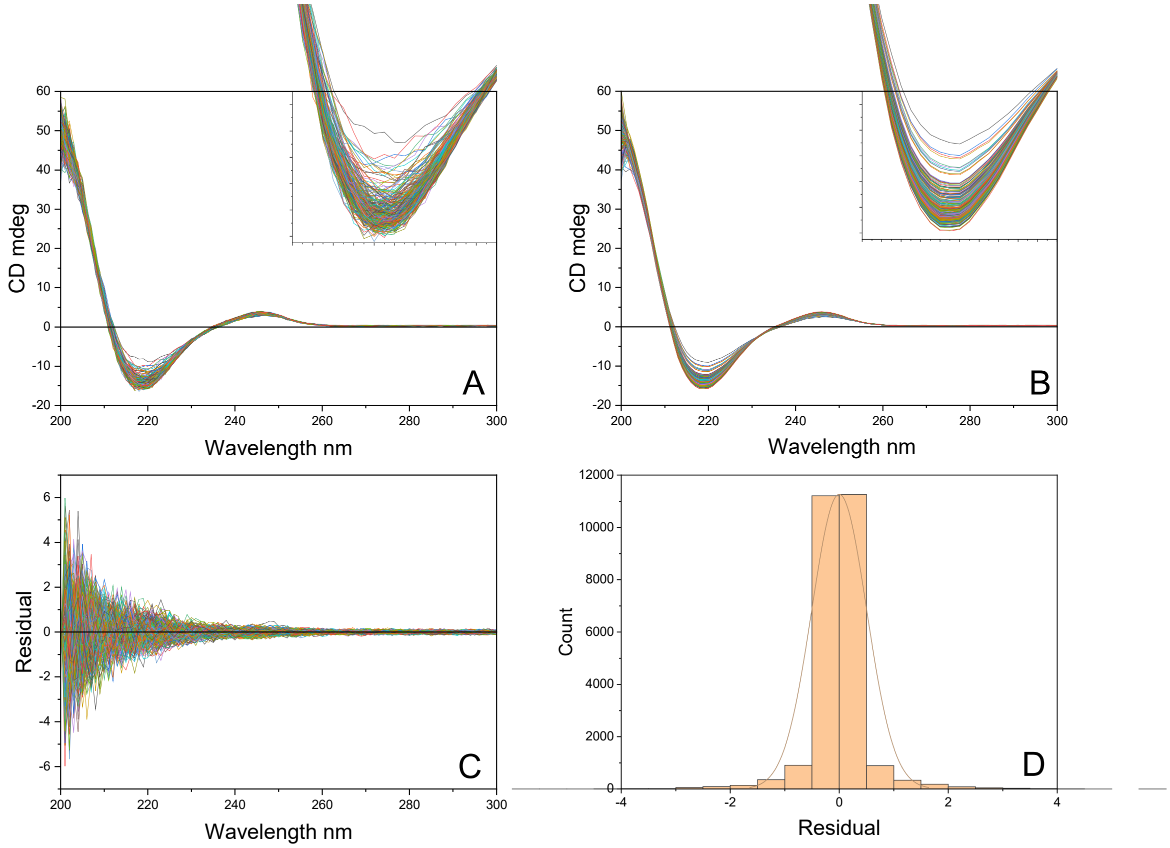


**Figure S2**. *Data reconstruction from SVD*. To validate the conclusion that the temporal evolution of the spectral changes upon assembly arise from a two-step process with only two spectral species (and their difference), we reconstructed a data matrix $\mathbf{D_{calc}}$ using only the first 3 singular values. A diagonal matrix **S’** was constructed with all elements set to 0 except for the first 3 diagonal elements that were set to the values determined in the analysis of the primary data. The reconstruction then is $\mathbf{D_{calc}} = \mathbf{S'UV^T}$. The results are shown in panel B, compared to the original primary data in panel A. As is best seen in the insets, the shapes and temporal evolution of the spectra are retained in the reconstruction, but noise has been removed. The differences in the primary and reconstructed data are superimposed and shown in panel C. To be clear, these differences correspond to the difference between an indivual spectrum at a given time minus its reconstructed counterpart, $\mathbf{D} - \mathbf{D_{calc}}$. The distribution of these residuals is shown in panel D. The residuals from the 3-component SVD reconstruction are centered essentially at zero (mean $\approx -8 \times 10^{-5}$) with a standard deviation of $\approx 0.50$ over a range of roughly -6 to +6 CD units, indicating an unbiased fit with most points closely reproduced. The distribution is nearly symmetric (skewness $\approx$ -0.04) but extremely leptokurtic (kurtosis $\approx$ 22), reflecting a very sharp peak at zero accompanied by heavy tails arising from a relatively small number of large residuals. Quantitatively, this behavior is consistent with heteroscedastic errors dominated by noisy low-wavelength CD data, while the first three singular values, which together account for $\approx$ 99.8% of the variance, adequately capture the underlying spectral structure.

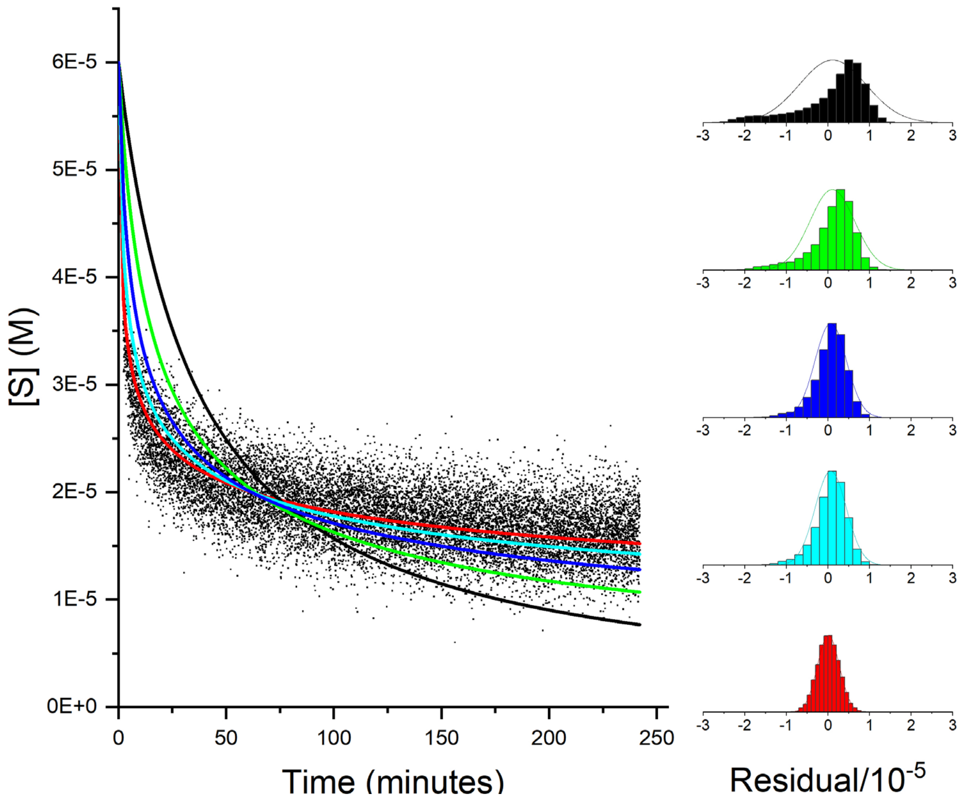


**Figure S3.** *Determination of reaction order*. Using representative kinetic data with 60 μM total strand concentration the apparent reaction order was determined. The reaction order is defined as $\frac{dS}{dt} = -nk[S]^n$. The kinetic curve was fit to that equation for values on n ranging from 2 to 6 using the "Fit ODE" app in Origin software. The fitted curves are shown from black (n=2) to red (n=6), and the distribution of residuals from the fits (scaled to identical y-axis limits) are shown on the right with the same color scheme. The magnitudes of the residuals decreases with increasing values of n, and the shape of the residuals plots evolve from skewed (n=2) to Gaussian randomness at n=6. These residual plots show clearly that the optimal fit is with n=6, meaning that the assemble reaction follows a rare apparent sixth-order law. Since all reactants are not consumed, the reaction equilibrates by 240 minutes, indicating that the assembly reaction is reversible with a necessary contribution from a reverse rate constant.

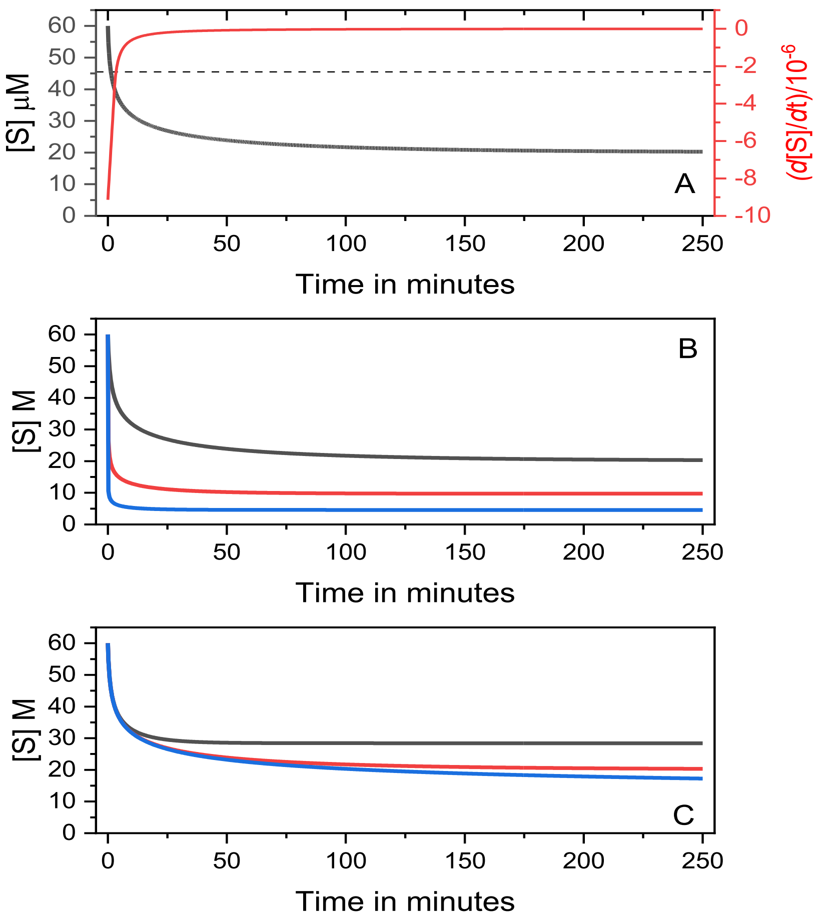


**Figure S4**. *Simulations of a reversible sixth-order reaction*. A) Simulated kinetics for 60 μM total strand concentration with $k_f = 10^{20}$ $M^{-5}$ $min^{-1}$ and $k_r = 0.001$ $min^{-1}$ (black). The reaction half time is shown by the dashed line. The first derivative of that curve is shown in red. The reversible single-step sixth-order reaction is characterized by a sharp burst phase during which nearly half of the reactants are consumed within 2 min, followed by a slower crawl phase that equilibrates to an extent of reaction of 0.3. The red derivative curve emphasized the rapid initial rate and the full equilibration by 240 min. B) The effect of $k_f$ is shown in simulated data with 60 μM total strand, $k_r$=0.001. $k_f$ (in $M^{-5}$ $min^{-1}$) was set at $10^{20}$ (black), $10^{22}$ (red) and $10^{24}$ (blue). C) The effect of the reverse rate constant is shown with $k_f = 10^{20}$ $M^{-5}$ $min^{-1}$, 60 μM total strand and $k_r$ ($min^{-1}$) values of 0.01 (black), 0.001 (red) and 0.0001(blue). The simulations show that the forward rate constant affects both the initial rate of reaction and the final equilibrium position, while the reverse rate constant primarily affects the latter.

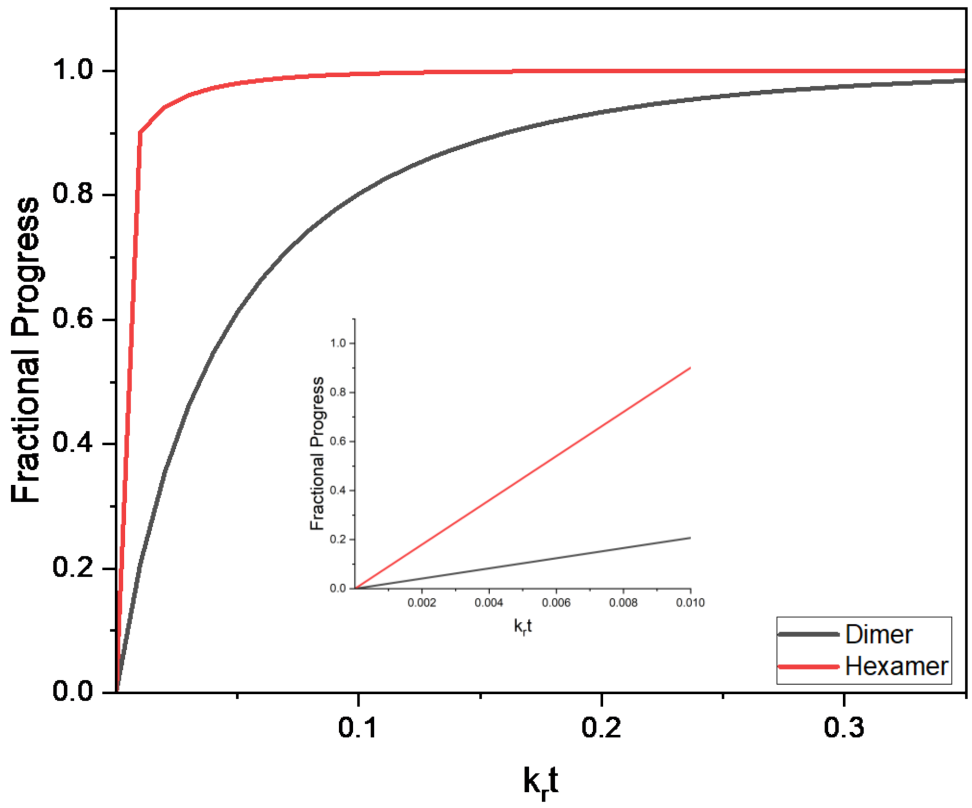


**Figure S5.** *Normalized kinetic progress curves for reversible dimer and hexamer assembly.* Simulations for $2S \rightleftharpoons S_2$ and $6S \rightleftharpoons S_6$ were constrained to the same equilibrium extent of assembly and used a common dissociation rate constant $k_r$. Fractional progress, $x/x_\infty$, is plotted versus dimensionless time, $\tau = k_r t$, where $x = [M_n]/C_0$. Forward rate constants were separately adjusted for each reaction order to yield the common equilibrium endpoint. The inset expands the early time range. The sixth-order model shows an exceptionally rapid but continuous initial burst, followed by a strongly concentration-sensitive approach to equilibrium.

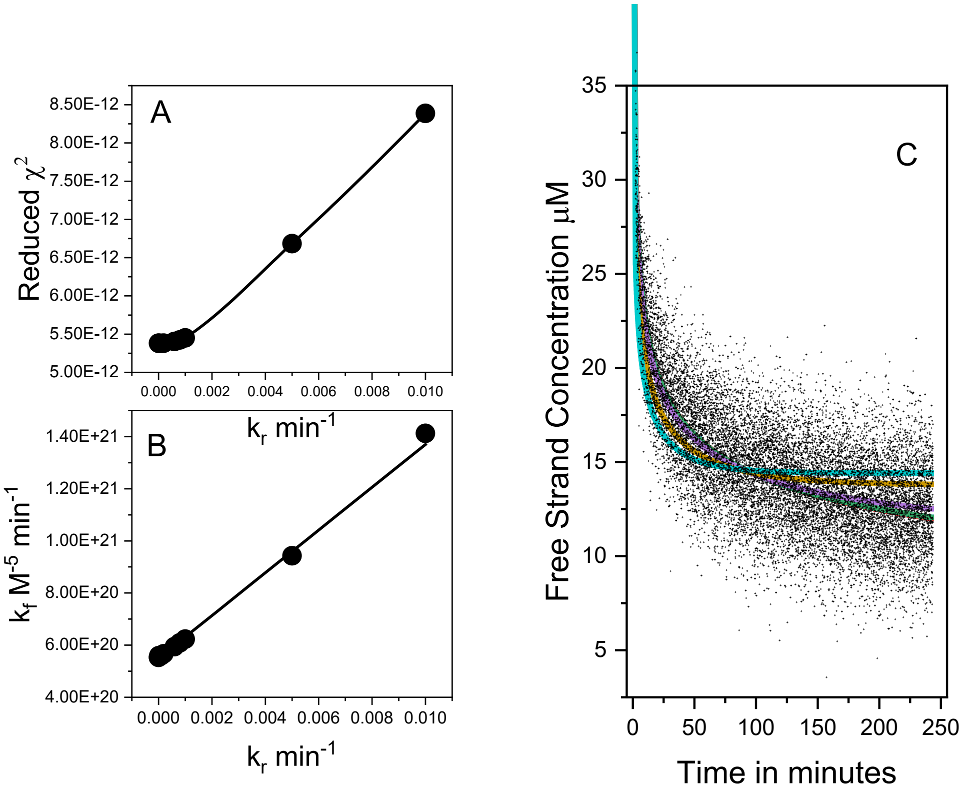


**Figure S6**. *Mapping the error space of $k_r$.* Primary kinetic data at 60 μM total strand concentration were fit with fixed values of the reverse rate constant $k_r$, while optimizing the forward rate constant $k_f$. Panel A shows the reduced $\chi^2$as a function of $k_r$; the reduced $\chi^2$remains nearly constant over the range 0 to $8 \times 10^{-4} min^{-1}$, indicating that this dataset provides little information to uniquely determine $k_r$and that all values in this interval fit the data essentially equally well. Panel B shows the corresponding fitted values of $k_f$, which vary systematically with $k_r$, revealing a strong correlation between the two rate constants and confirming that the inverse problem is ill-conditioned, no doubt because of the extreme disparity in parameter scales. Panel C compares best-fit kinetic curves obtained with fixed $k_r$values. The curves for $k_r = 0$ to 0.001 $\mathrm{min}^{-1}$ are essentially superimposed, whereas fits with larger $k_r$values (e.g., 0.005, red; 0.01, green) are clearly inadequate. Together, these results demonstrate that the 60 μM kinetic experiment robustly constrains $k_f$but renders $k_r$ non-identifiable over a broad range.

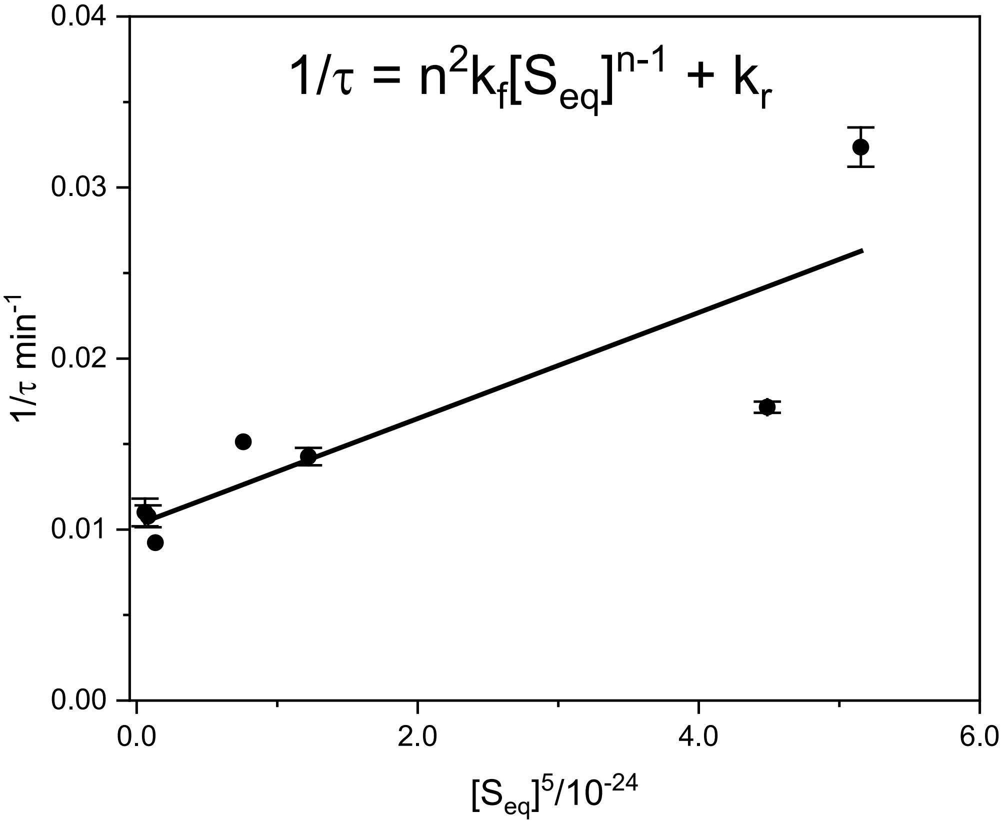


**Figure S7**. *Relaxation kinetic analysis of the reversible sixth-order assembly reaction*. The method of relaxation kinetics facilitates linearization of the differential equations governing rates of reactions, reducing the complexity and difficulty of the data fitting problem. Data here are cast into the linear form for analysis of a reversible sixth-order assembly mechanism using the relaxation time for the approach to equilibrium, with the linear closed-form fitting function shown at the top of the figure[5]. $S_{eq}$ is the concentration of free strands at equilibrium. The slope of the fitted line is $n^2 k_f$, from which a forward rate constant of $k_f = (8.6 \pm 0.2) \times 10^{19}$ M$^{-5}$ min$^{-1}$ where a reaction order of n=6 is assumed. The intercept is the reverse rate constant $k_r = 0.01 \pm 0.002$ min$^{-1}$. These rate constants yield an equilibrium constant $K_{eq} = \frac{k_f}{k_r} = 8.6 \times 10^{21}$ M$^{-5}$ min$^{-1}$ that agrees to within a factor of 2-3 with the value determined independently from the equilibrium analysis of the "hockey-stick" isotherm in Figure 2A.

**Table 1.** Optimized microscopic rate constants for sequential assembly model

| Reaction Step | $\mu M^{-1}min^{-1}$ | $M^{-1}min^{-1}$ |
|---|---|---|
| **kf1 (S + S -> D)** | 0.0268 | $2.68\times10^{04}$ |
| **kf2 (D + S -> T** | 1.9802 | $1.98\times10^{06}$ |
| **kf3 (T + S -> Q)** | 2.1499 | $2.15\times10^{06}$ |
| **kf4 (Q + S -> P)** | 2.1070 | $2.11\times10^{06}$ |
| **kf5 (P + S -> H)** | 0.2925 | $2.92\times10^{05}$ |

kr1 to kr4 (Intermediates Breakdown) : 100.00 $min^{-1}$

kr5 (Hexamer Reversible Sink Rate) : $1.15\times10^{-4}$ $min^{-1}$

Calculated Apparent Macroscopic kf = $7.0215\times10^{20}$ $M^{-5}$ $min^{-1}$
